# ssc-micorRNA-132 suppresses the *Clostridium perfringens* beta2 toxin induced inflammation and apoptosis of IPEC-J2 cells via targeting *DACH1*

**DOI:** 10.1101/2020.11.24.396697

**Authors:** Kaihui Xie, Zunqiang Yan, Wei wang, Ruirui Luo, Xiaoli Gao, Pengfei Wang, Qiaoli Yang, Xiaoyu Huang, Juanli Zhang, Jiaojiao Yang, Shuangbao Gun

**Affiliations:** College of Animal Science and Technology, Gansu Agricultural University, Lanzhou Gansu China 730070; Gansu Research Center for Swine Production Engineering and Technology, Lanzhou Gansu China 730070

**Author notes:** Corresponding author: Shuangbao Gun.

**Keywords:** ssc-miR-132, recombinant *Clostridium perfringens* beta2 (rCPB2) toxin, IPEC-J2 cells, *DACH1*, apoptosis, inflammation

## Abstract

*Clostridium perfringens* (*C. perfringens*) beta2 (CPB2) is the main virulence factor secreted from *C. perfringens* type C, which caused diarrhea characterized by high mortality in pig, especially newborn piglets. Our previous research found that ssc-miR-132 displayed decreased expression in piglets diarrhea after infected with *C. perfringens* type C compared with normal piglets. We speculated that ssc-miR-132 may play an important role in the diarrhea. However, the function of ssc-miR-132 in the diarrhea is limited. Thus, we overexpressed and knocked down ssc-miR-132 in intestinal porcine epithelial (IPEC-J2) cells, and then treated the cells with recombinant CPB2 (rCPB2) toxin (20 μg/mL). Our results showed that ssc-miR-132 was significantly decreased after treated with rCPB2 toxin. In addition, overexpression of ssc-miR-132 reduced the expression of lactate dehydrogenase (LDH) and tumor necrosis factor (*TNF-α*), interleukin-6 (*IL-6*) and interleukin-8 (*IL-8*) caused by rCPB2 toxin. The CCK8, Edu and TUNEL staining showed that overexpression of ssc-miR-132 weakened the inhibition of rCPB2 toxin on cell proliferation and reduced the promotion of cell apoptosis; while inhibition of ssc-miR-132 had opposite results. The dual luciferase experiment showed that dachshund family transcription factor 1 (*DACH1*) was the target gene of ssc-miR-132. Silencing *DACH1* was consistent with the results of overexpression of ssc-miR-132, and reversed the apoptosis and inflammation caused by rCPB2 toxin. Overexpression of *DACH1* weakened the role of ssc-miR-132 in rCPB2 toxin -induced inflammation and apoptosis. In summary, ssc-miR-132 inhibited rCPB2 toxin-induced apoptosis and inflammation in IPEC-J2 cells by targeting *DACH1*.

## Introduction

*Clostridium perfringens* (*C. perfringens*) can cause intestinal infections and tissue toxic diseases in animals (1, 2).It is classified into types A to E according to their ability to produce types of toxins (3). *C. perfringens* type C is one of the main pathogens leading to piglet diarrhea (4), and its infection caused piglet mortality rate as high as 70%, which brought serious losses to the pig industry (5). *C. perfringens* beta2 (CPB2) toxin is the main virulence factor of *C. perfringens* type C and causes a severe necrotic and hemorrhagic enteritis in pig, sheep, cattle, human and so on (6, 7). Pigs are most easily infected with *C. perfringens* type C, and the necrotic enteritis caused by it causes high mortality in newborn piglets (8). CPB2 toxin can cause intestinal mucosal damage and be toxic to pig intestinal epithelial cells (9, 10). However, there is little information on the specific molecular mechanism of CPB2 toxin induced intestinal inflammatory damage. Therefore, studying its mechanism of action in intestinal porcine epithelial (IPEC-J2) cells will help the prevention and control of piglet diarrhea.

MicroRNAs (miRNAs) are endogenous 19-23 nucleotide (nt) RNAs and pair with mRNAs of protein-coding genes to suppress translation and degrade mRNAs in post-transcriptional level (11). miRNAs are related to cellular proliferation, differentiation, apoptosis, migration, invasion and inflammation (12–16). Additionally, miRNAs play a crucial regulatory role in host immune response to pathogenic infection. For example, gga-miR-200a-3p regulated chicken necrotic enteritis caused by *C. perfringens* through the regulation of MAPK signaling pathway related genes *TGFβ2*, *ZAK* and *MAP2K4* (17). Ye et al. (18) discovered a total of 11 up-regulated expression miRNAs and 1 down-regulated miRNA associated with *Escherichia coli* (ETEC) F18 infection susceptible piglet diarrhea. Zhang et al (19) found that miR-128 regulated Salmonella typhimurium infection by targetng *M-CSF*. Hoeke et al. (20) observed that miR-29a-mediated caveolin-2 regulated intestinal epithelial cell proliferation and Salmonella uptake. miR-143 targeted *ATP6V1A* and *IL13RA1*, miR-26 targeted *BINP3L* and *ARL6IP6* to regulate Salmonella infection in pigs (21).

miRNA-132 is a multifunctional miRNA. miR-132 is closely related to inflammation, and plays an important role in viral infections, bacterial infections, wound healing, and inflammation induced by nutritional deficiency (22–26). Liu et al. (27) showed that miR-132-3p reduced LPS-induced inflammation by inhibiting the NF-κB pathway activated by *FOXO3*. Shaked et al. (28) found that miR-132 targeted acetylcholinesterase (AchE) and reduced the hydrolysis of acetylcholine (Ach), achieved a cholinergic effect, regulated the choline pathway, and inhibited inflammation in mice. However, the functions of ssc-miR-132 in intestinal inflammation caused by *C. perfringens* type C infection is unclear.

This study aimed to determine the role of ssc-miR-132 in rCPB2-induced inflammation of IPEC-J2 cells by evaluating cell viability, apoptosis and inflammatory damage. This study provides a reference for the prevention and treatment of piglet diarrhea.

## Results

### Conserved ssc-miR-132 was down-regulated in rCPB2-induced IPEC-J2 cells

Through sequence alignment, the miR-132 mature sequence was conserved in many species (such as humans, pigs and cattle, etc) via alignment, which indicated that conservative miR-132 play important role in biological (Fig. 1A). In previous work, ssc-miR-132 was down-regulated in diarrheic piglets infected with *C. perfringens* type C (susceptible group, IS) compared with the control group (IC) (Fig. 1B). The IPEC-J2 cells were treated with rCPB2 toxin for 24 h, ssc-miR-132 expression level was significantly down-regulated (Fig. 1C). ssc-miR-132 mimics, inhibitor and their respective negative controls were utilized to transfect IPEC-J2 cells. We found that ssc-miR-132 was significantly up-regulated after transfection of mimics, and it was down-regulated after transfection of inhibitor (Fig. 1D).

**Figure 1.**
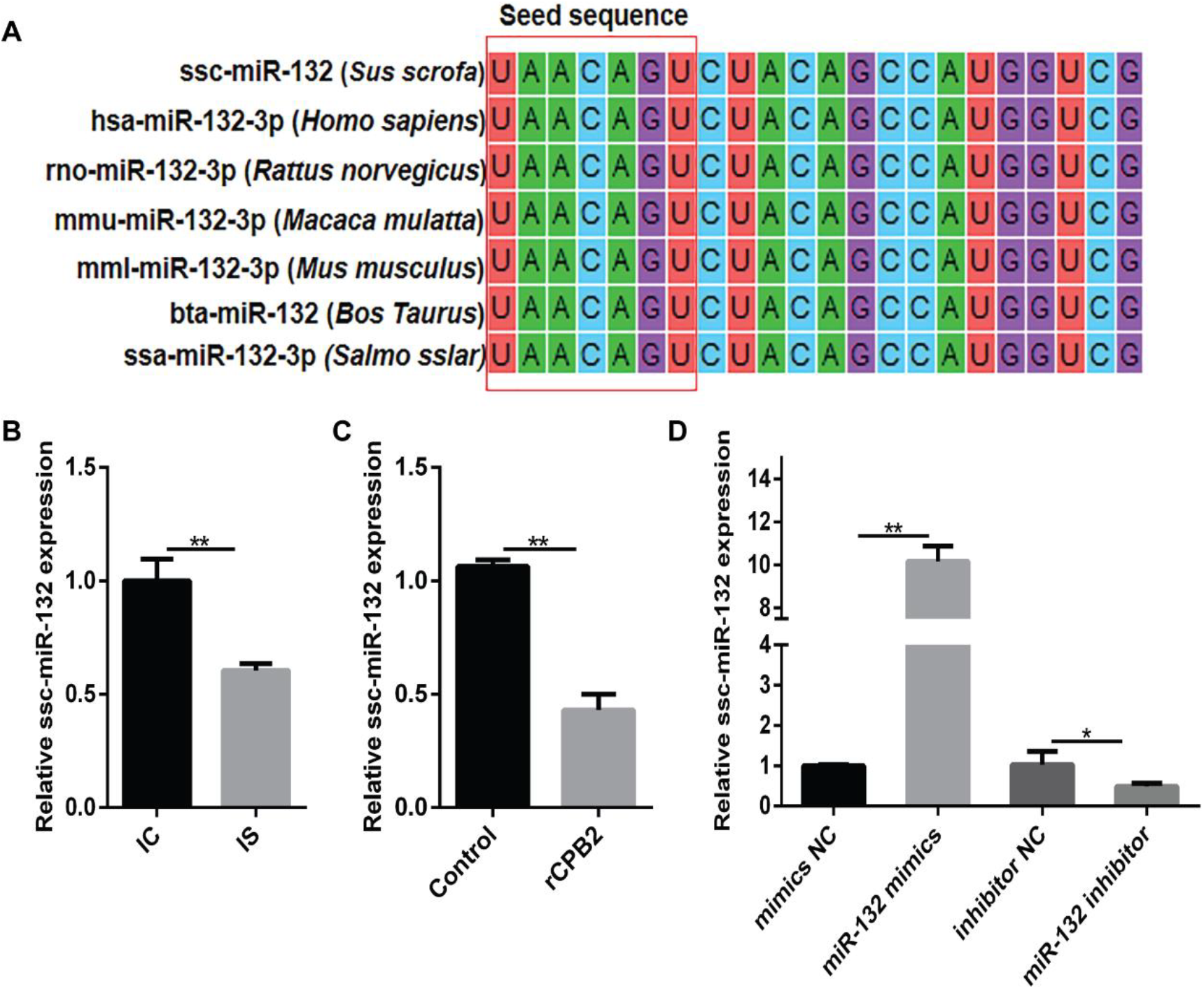
Conserved ssc-miR-132 was down-regulated in rCPB2-induced IPEC-J2 cells. **A)** miR-132 was highly conserved among different species. **B)** ssc-miR-132 was down-regulated in diarrheic piglets infected with *C. perfringens* type C. **C)** ssc-miR-132 was down-regulated in rCPB2 toxin-treated IPEC-J2 cells. **D)** The expression of ssc-miR-132 after transfection of miR-132 mimics, inhibitor and their respective negative controls in IPEC-J2 cells. * *P* < 0.05, ** *P* < 0.01.

### rCPB2-induced inflammation of IPEC-J2 cells was ameliorated by overexpression of ssc-miR-132

To verify whether ssc-miR-132 was involved in rCPB2 toxin-induced inflammatory response in IPEC-J2 cells, ssc-miR-132 mimics, inhibitor and their respective negative controls were transfected into cells. The transfected cells were treated with rCPB2 for 24 h. Lactate dehydrogenase (LDH) assay results showed that rCPB2 promoted the release of LDH, while overexpression of ssc-miR-132 reduced the release of LDH caused by rCPB2 (Fig. 2A). The expression of inflammatory cytokines was detected by qRT-PCR. rCPB2 facilitated the expression of tumor necrosis factor (*TNF-α*), interleukin 6 (*IL-6*) and interleukin 8 (*IL-8*), however ssc-miR-132 overexpression suppressed three cytokines expression (Fig. 2B-D). The expression level of ssc-miR-132 was opposite to the expression of inflammatory cytokines.

**Figure 2.**
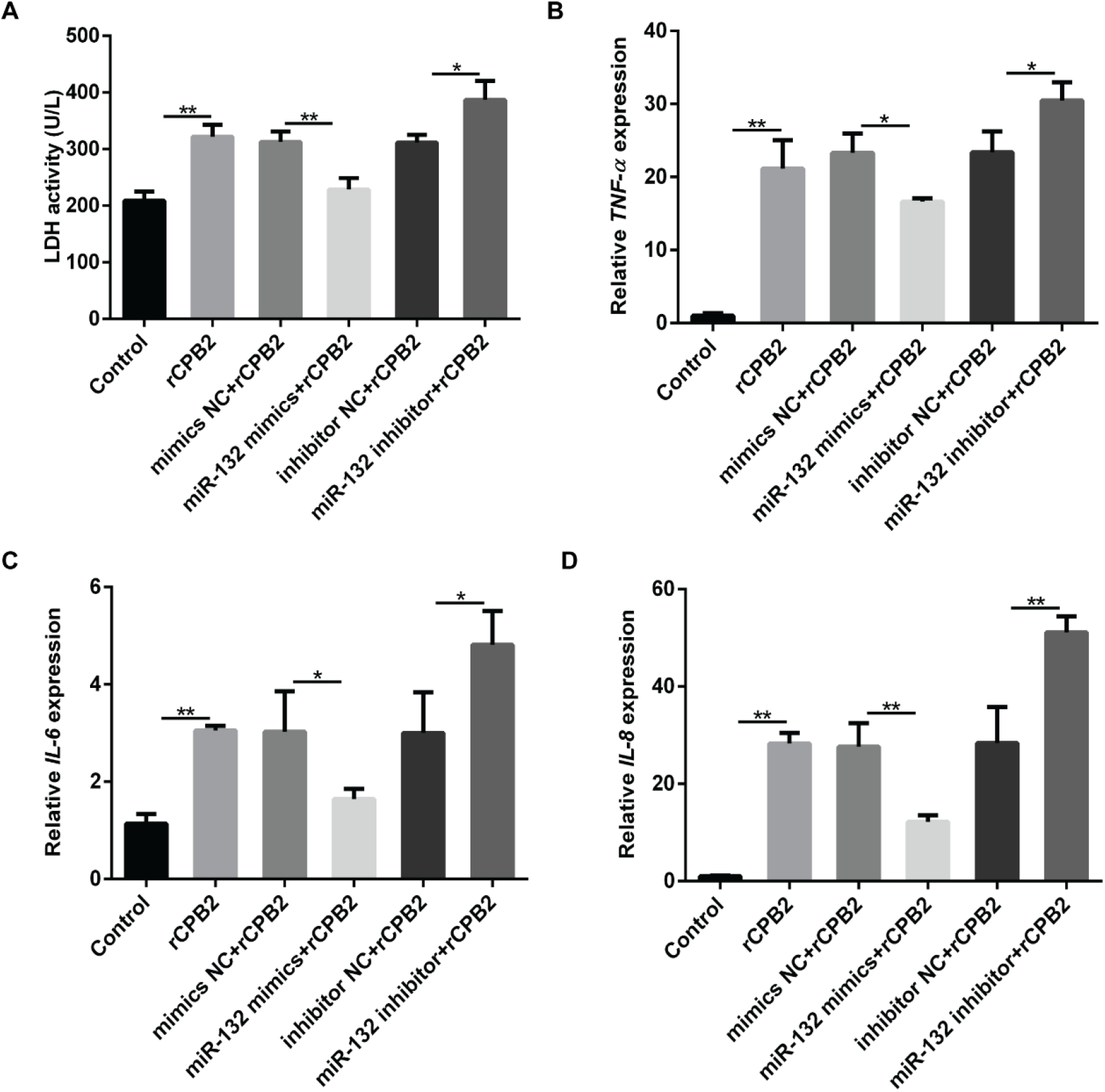
ssc-miR-132 attenuated the cytotoxicity and inflammation caused by rCPB2. **A)** Overexpression of ssc-miR-132 attenuated the cytotoxicity caused by rCPB2. **B-D)** Up-regulation of ssc-miR-132 suppressed rCPB2-induced release of pro-inflammatory factors (*TNF-α*, *IL-6* and *IL-8*). * *P* < 0.05, ** *P* < 0.01.

### Overexpression of ssc-miR-132 attenuated inhibition of cell viability and promoted cell proliferation caused by rCPB2

IPEC-J2 cells were transfected with ssc-miR-132 mimics, inhibitor and their respective negative controls to investigate the functions of ssc-miR-132 in rCPB2-affected IPEC-J2 cell proliferation. rCPB2 reduced the cell survival rate, while ssc-miR-132 mimics alleviated this inhibition, and ssc-miR-132 inhibitor promoted the inhibition of cell viability by rCPB2 (Fig 3A). The Edu staining also showed the same results. rCPB2 suppressed cell proliferation, but ssc-miR-132 mimics weakened this inhibition (Fig 3B-C).

**Figure 3.**
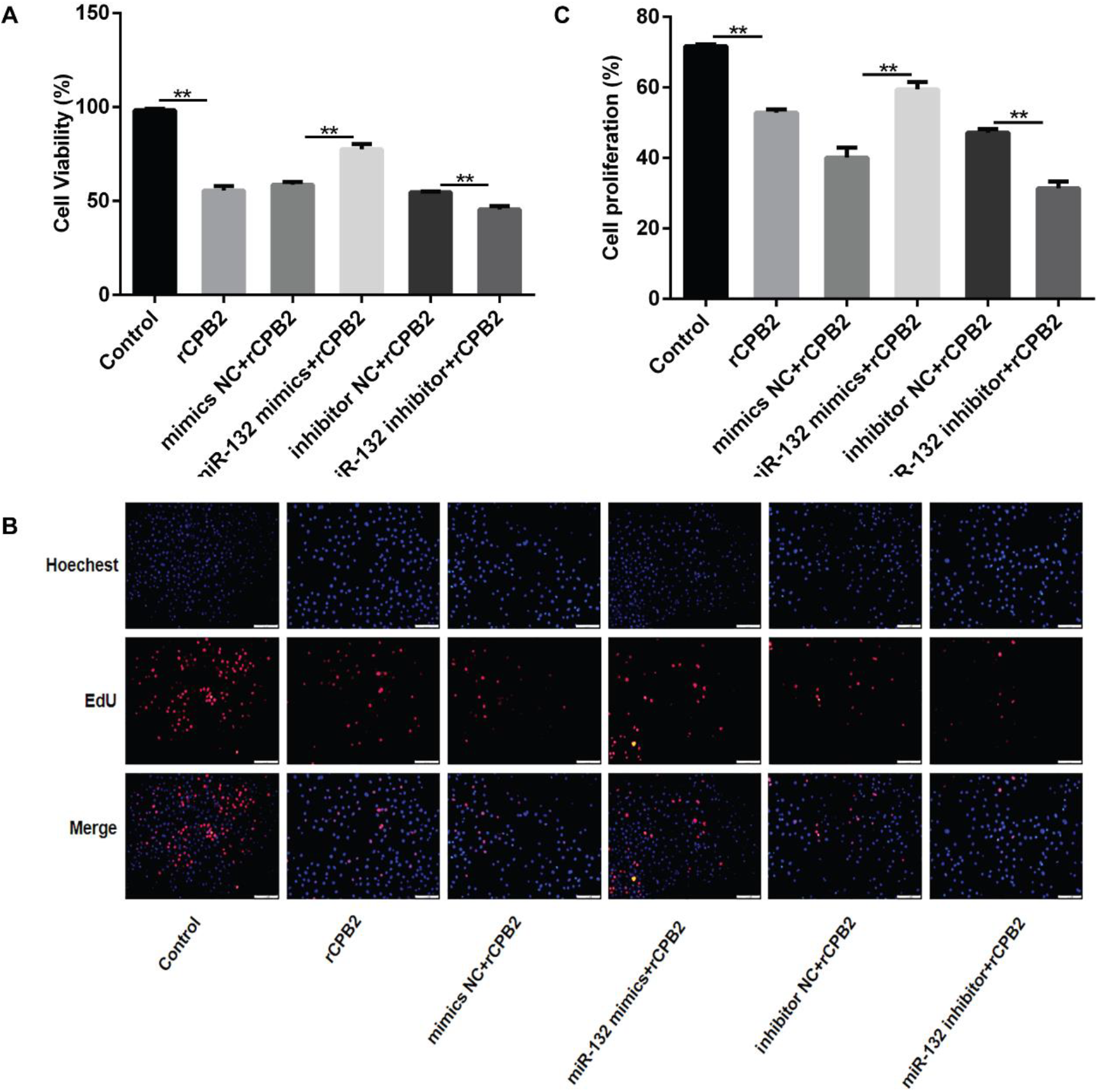
Overexpression of ssc-miR-132 reversed the inhibition of rCPB2 on cell proliferation. **A)** Up-regulation of ssc-miR-132 alleviated the suppression of rCPB2 on cell viability. **B-C)** Overexpression of ssc-miR-132 weakened the inhibition of rCPB2 on cell proliferation, and knockdown of ssc-miR-132 promoted the inhibition of rCPB2 on cell proliferation. ** *P* < 0.01

### Up-regulated ssc-miR-132 reduced the rCPB2-induced cell apoptosis

The TUNEL staining was used to verify the role of ssc-miR-132 in rCPB2-induced cell apoptosis. After ssc-miR-132 mimics, inhibitor and their respective negative controls transfected IPEC-J2 cells, cells were treated with rCPB2 for 24 h. The result showed that rCPB2 accelerated the apoptosis of IPEC-J2 cells, while ssc-miR-132 mimics significantly inhibited the cell apoptosis, and ssc-miR-132 inhibitor promoted the apoptosis caused by rCPB2 (Fig 4A-B). The Western Blot results showed that rCPB2 promoted the expression of Bax and Caspase3 protein and inhibited the expression of Bcl-2, while ssc-miR-132 mimics weakened the expression of Bax and Caspase3 and promoted the expression of Bcl-2. (Fig 4C-D).

**Figure 4.**
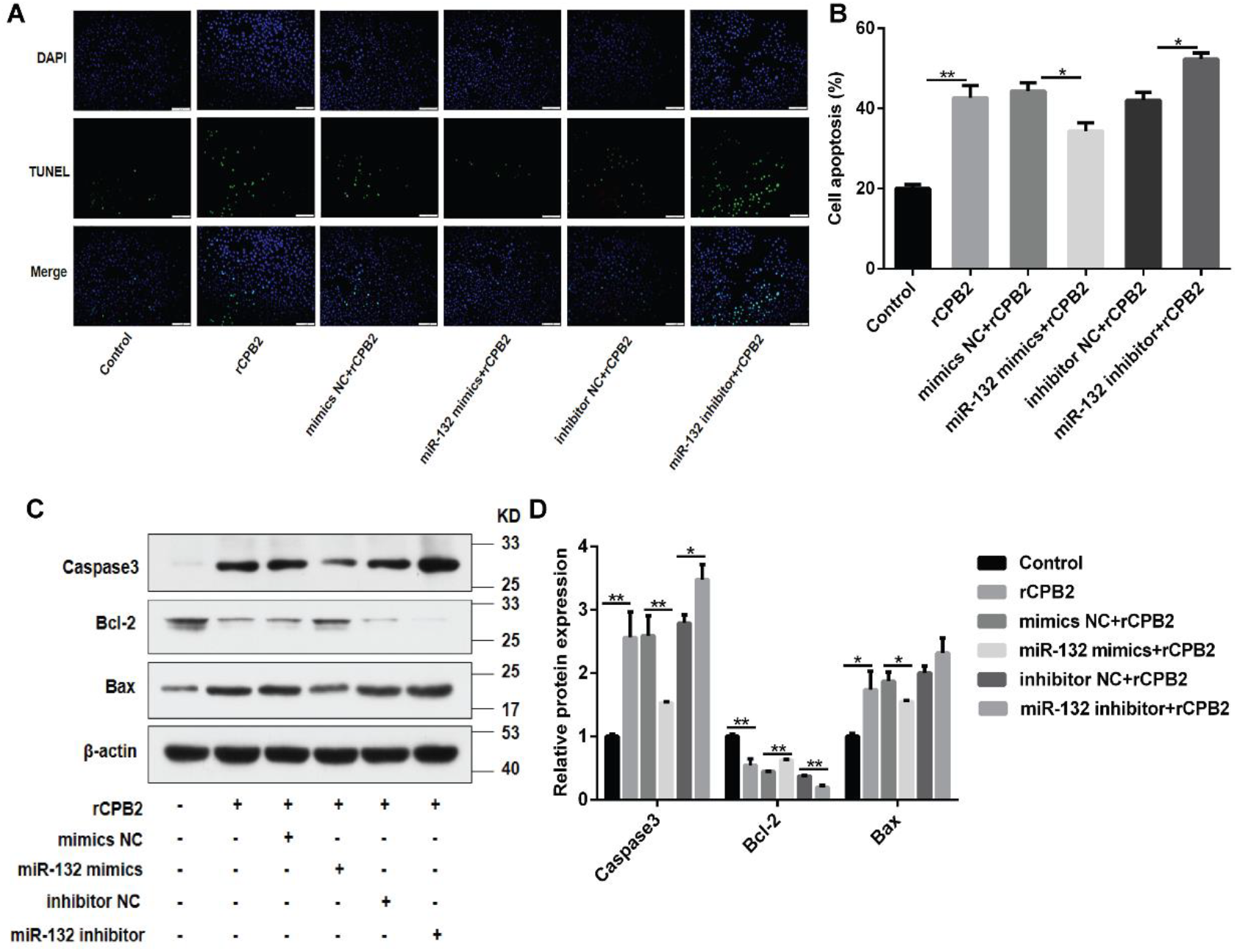
Up-regulation of ssc-miR-132 alleviated rCPB2-induced apoptosis. **A-B)** Overexpression of ssc-miR-132 reduced rCPB2-induced apoptosis, and knockdown aggravated rCPB2-induced apoptosis. **B)** Expression of pro-apoptotic proteins Caspase3, Bax and anti-apoptotic protein Bcl-2,after overexpression and knockdown of ssc-miR-132. * *P* < 0.05, ** *P* < 0.01.

### *DACH1* was the target gene of ssc-miR-132

Through PicTar, miRDB and TargetScan, we predicted that *DACH1* was the candidated target gene of ssc-miR-132 (Fig 5A). We verified that *DACH1* was up-regulated in rCPB2-treated IPEC-J2 cells, which was contrary to the expression of ssc-miR-132 (Fig 5B). Furthermore, after overexpression of ssc-miR-132, the DACH1 expression was decreased at both the mRNA and protein levels (Fig 5C-E). To determine whether ssc-miR-132 directly acts on *DACH1*, we constructed pmirGLO-DACH1 3’UTR-WT and pmirGLO-DACH1 3’UTR-Mut vectors (Fig 5F). Luciferase reporter gene assay showed that when pmirGLO-DACH1 3’UTR-WT was co-transfected with ssc-miR-132 mimics, relative luciferase activity was significantly reduced compared with negative controls. Whereas there was no effect in luciferase activity from the mutated reporter vector (Fig 5G). The above results indicated that *DACH1* was the target gene of ssc-miR-132.

**Figure 5.**
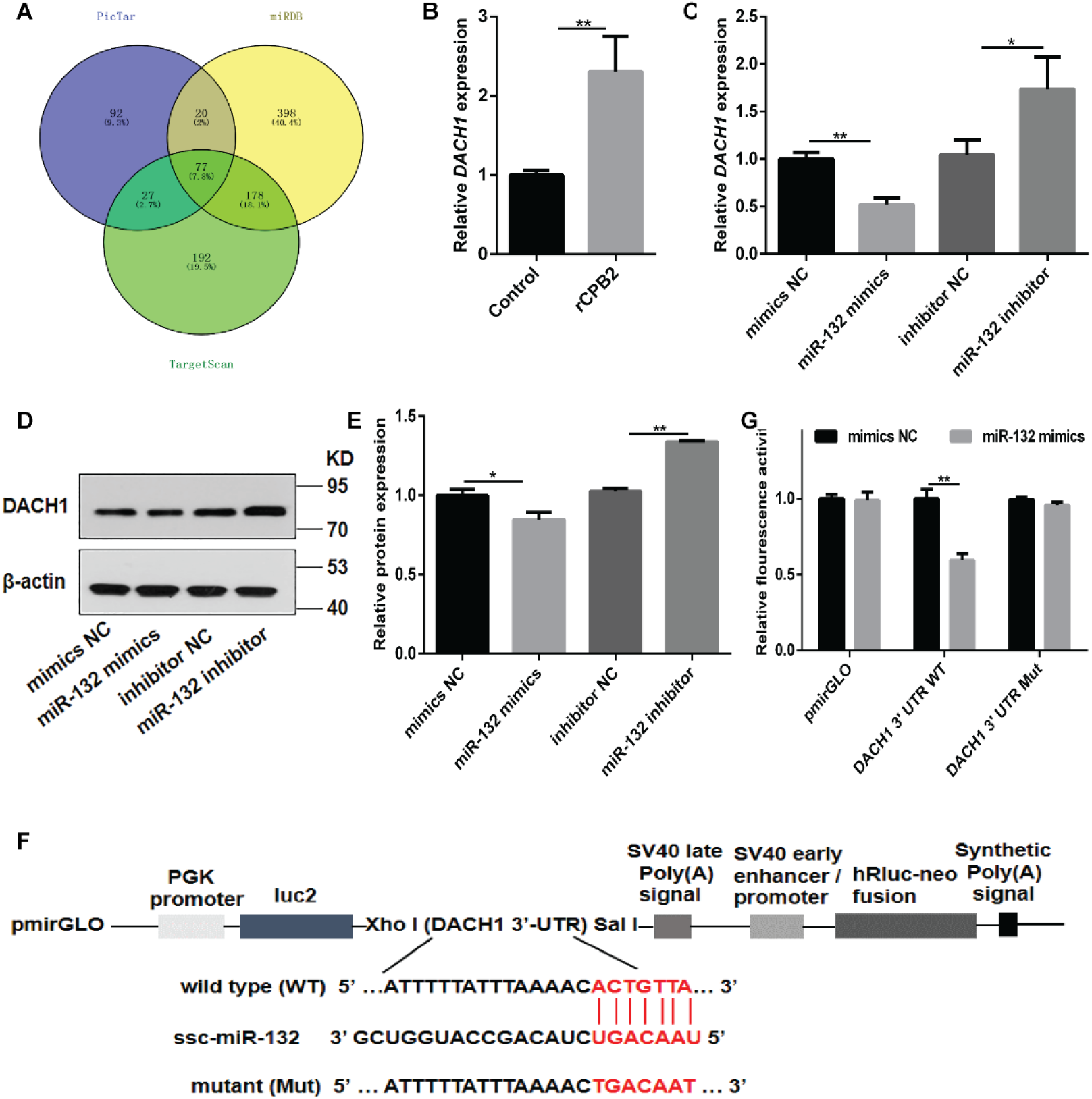
*DACH1* was the target gene of ssc-miR-132. **A)** ssc-miR-132 target gene prediction. **B)** *DACH1* was down-regulated in rCPB2-treated IPEC-J2 cells. **C)** Expression of *DACH1* after overexpression and knockdown of ssc-miR-132. **D-E)** Expression of DACH1 protein after overexpression and knockdown of sc-mir-132. **F)** Construction diagram of dual luciferase reporter vector containing wild-type and mutant DACH1 3’-UTR sequences. **G)** Fluorescence activity of 293T cells after the wild-type and mutant vectors of *DACH1* were cotransformed with miR-132 and mimics NC, respectively. * *P* < 0.05, ** *P* < 0.01.

### Silencing *DACH1* inhibited the expression of pro-inflammatory cytokines

In order to verify the function of *DACH1* in rCPB2-induced IPEC-J2 cells, cells were transfected with pcDNA3.1-DACH1, si-DACH1 and their respective negative controls (pcDNA3.1 vector and si-NC). The results showed that DACH1 was successfully overexpressed and silenced at the mRNA and protein levels (Figure 6A-C). Then IPEC-J2 cells were treated with rCPB2 toxin for 24 h. Down-regulation of *DACH1* attenuated the release of LDH and the expression of *TNF-α*, *IL-6* and *IL-8* caused by rCPB2 (Fig 6D-G). The above results indicated that knockdown of *DACH1* alleviated the inflammatory response induced by rCPB2.

**Figure 6.**
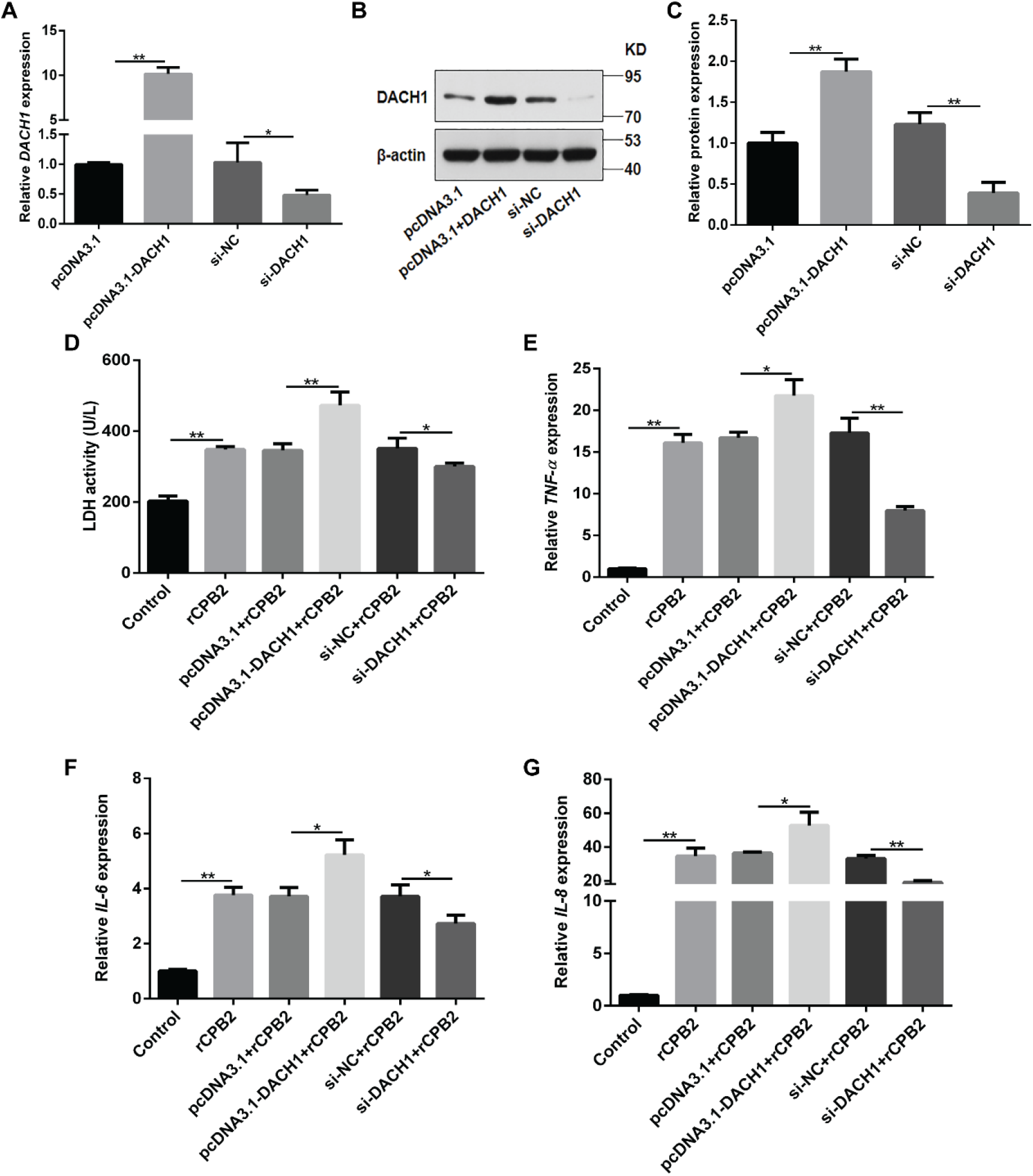
Down-regulation of *DACH1* alleviated rCPB2-induced inflammation. **A)** The expression of *DACH1* after transfection of pcDNA3.1-DACH1, si-DACH1 and their respective negative controls. **B-C)** The expression of DACH1 protein after overexpression and knockdown of DACH1. **D)** Knockdown of *DACH1* reduced rCPB2-induced cytotoxicity. **E-G)** Silencing *DACH1* alleviated the expression of pro-inflammatory factors (*TNF-α*, *IL-6* and *IL-8*) rCPB2-induced, while overexpression of *DACH1* promoted their expression. * *P* < 0.05, ** *P* < 0.01.

### Knockdown of *DACH1* promoted cell proliferation and lightened rCPB2-induced apoptosis

The IPEC-J2 cells were transfected with pcDNA3.1-DACH1, si-DACH1, pcDNA3.1 vector and si-NC and treated with rCPB2 for 24 h to determine the effect of *DACH1* on the proliferation and apoptosis of IPEC-J2 cells. The CCK8 assay indicated that down-regulated *DACH1* alleviated the inhibition of cell viability by rCPB2, and overexpression of *DACH1* is on the contrary (Fig 7A). The Edu staining also showed that silencing *DACH1* facilitated cell proliferation(Fig 7B-C). The TUNEL staining revealed that si-DACH1 alleviated the rCPB2-induced apoptosis, but pcDNA3.1-DACH1 promoted this apoptosis(Fig 7D-7E). Down-regulated *DACH1* inhibited the expression of Bax and Caspase3 proteins and promoted the expression of Bcl-2 (Fig 7F-7G).

**Figure 7.**
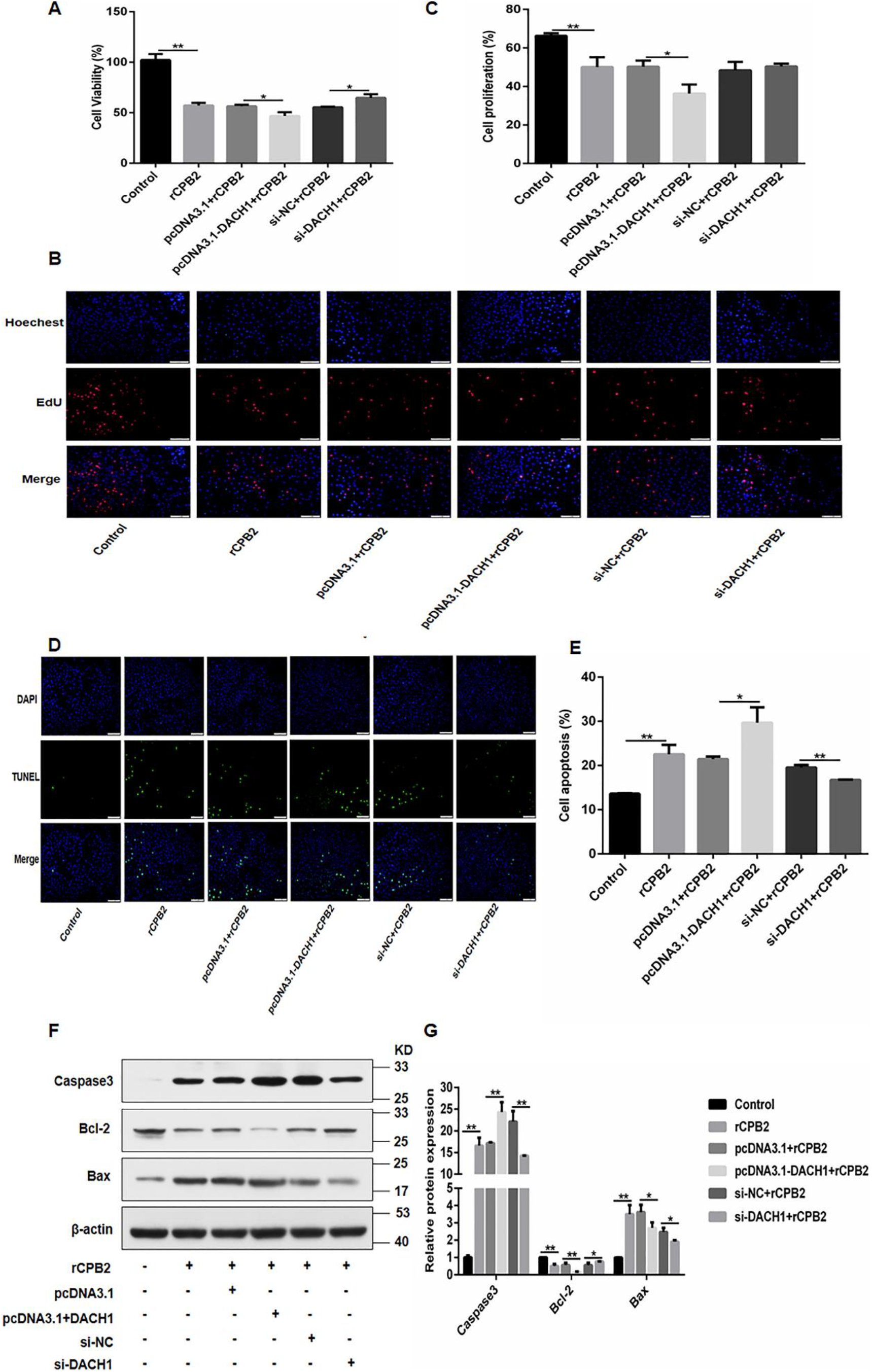
Silencing *DACH1* alleviated the suppression of rCPB2 on cell proliferation and attenuated rCPB2-induced apoptosis. **A)** Down-regulation of *DACH1* alleviated the inhibition of rCPB2 on cell viability. **B-C)** Silencing *DACH1* weakened the inhibitory effect of rCPB2 on cell proliferation. **D-E)** si-DACH1 inhibited rCPB2-induced cell apoptosis. **F-G)** The effect of silencing and overexpression of *DACH1* on apoptosis-related proteins. * *P* < 0.05, ** *P* < 0.01.

### Rescue experiment

We co-transfected ssc-miR-132 mimics and pcDNA3.1, ssc-miR-132 mimics and pcDNA3.1-DACH1 into IPEC-J2 cells and treated them with rCPB2 for 24 h to verify whether miR-132 exerted its functions through *DACH1* in rCPB2-induced inflammation. The results showed that compared with the co-transfection of miR-132 mimics and pcDNA3.1 group after co-transfection of miR-132 mimics and pcDNA3.1-DACH1 group, the expression of LDH, *TNF-α*, *IL-6* and *IL-8* were increased, and the cells viability was decreased (Fig 8A-C). These results indicated that *DACH1* reversed the resistance of ssc-miR-132 inflammation caused by rCPB2. The Edu assay demonstated that when miR-132 mimics and pcDNA3.1-DACH1 were co-transfected, the number of proliferating cells were less than that of the miR-132 mimics and pcDNA3.1 group (Fig 8D-E). The TUNEL staining indicated that *DACH1* attenuated the inhibitory effect of ssc-miR-132 on apoptosis (Fig 8F-8G). Compared with the co-transformed miR-132 mimics and pcDNA3.1 group, the expression of pro-apoptotic proteins Bax and Caspase3 increased in the co-transformed miR-132 mimics and pcDNA3.1-DACH1 group, while anti-apoptotic Bcl-2 was down-regulated (Fig 8H-8I). The above results suggested that ssc-miR-132 indeed resisted apoptosis and inflammation caused by rCPB2 through *DACH1*.

**Figure 8.**
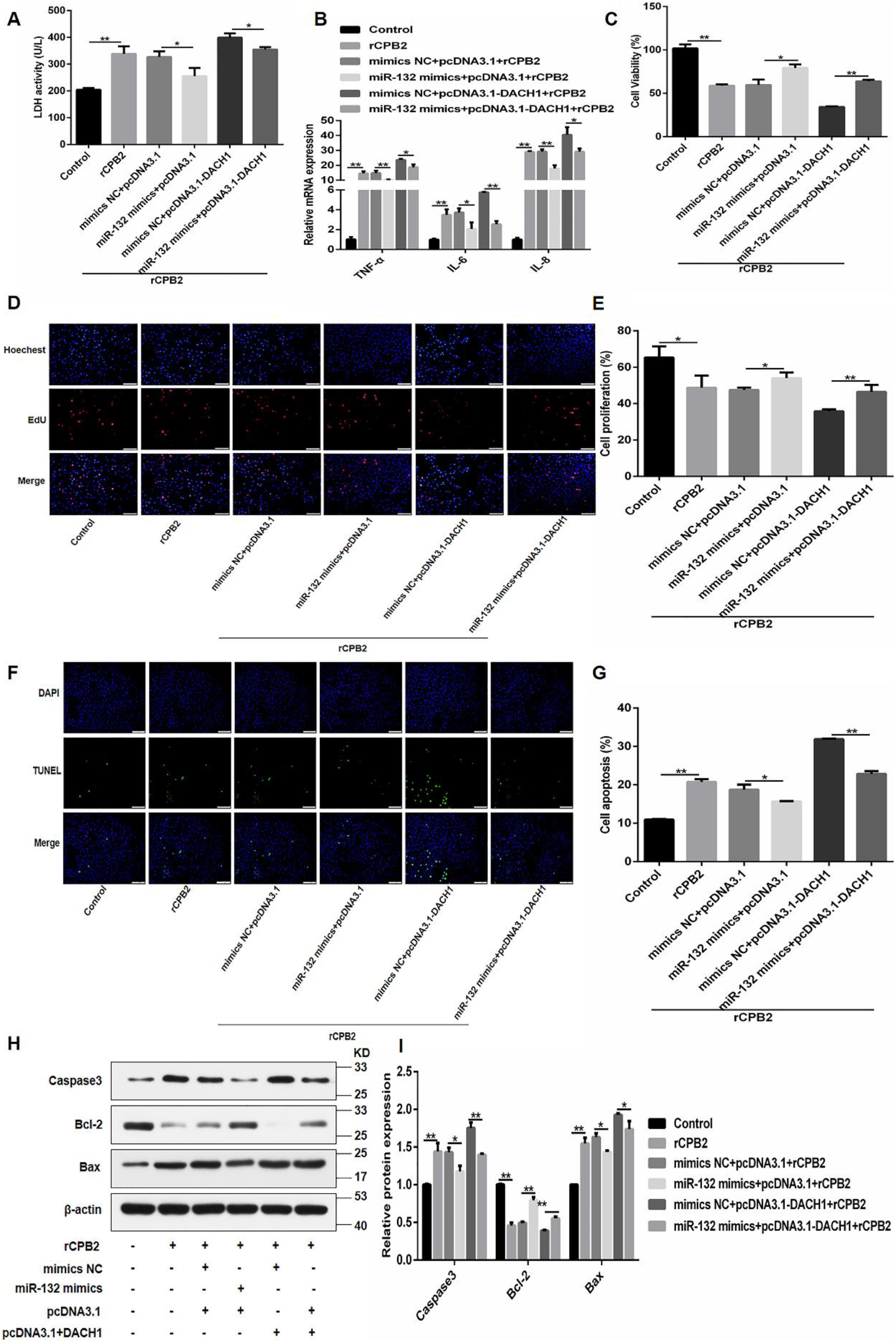
Overexpression of *DACH1* attenuated the role of ssc-miR-132 in rCPB2-induced inflammation. **A)** LDH activity. **B)** Proinflammatory cytokine expression. **C)** The effect of simultaneous overexpression of miR-132 mimics and *DACH1* on cell viability. **D-E)** Overexpression of *DACH1* attenuated the promoting effect of ssc-miR-132 on cell proliferation. **F-G)** pcDNA3.1-DACH1 reversed the inhibitory effect of ssc-miR-132 on cell apoptosis. **H-I)** up-regulated *DACH1* weakened the inhibitory effect of ssc-miR-132 on pro-apoptotic proteins and reversed the promotion of anti-apoptotic proteins. * *P* < 0.05, ** *P* < 0.01.

## Discussion

Diarrhea is a common infectious intestinal disease that causes the death of piglets. It has caused great economic losses to the pig industry and seriously hindered the healthy development of the pig industry. *C. perfringens* type C is one of the common pathogens causing bacterial diarrhea in piglets (4). *C. perfringens* type C can produce alpha and beta toxins (29). Beta toxin is the key toxin causing disease of *C. perfringens* type C (6). Beta toxin can cause intestinal vascular damage, hemorrhage and decreased oxygen content, causing tissue necrosis and promote the secretion of TNF-α and IL1β activates the Tachykinin NK1 receptor, causing inflammation in the animal body (30–32). CPB2 toxin cause intestinal wall bleeding and small intestinal loop necrosis (33, 34). The intestinal barrier can effectively resist external damage to the host, and its integrity has a protective effect on the body. Intestinal epithelial cells are an important part of the intestinal barrier, protecting the intestine from harmful substances such as bacteria, bacterial toxins and bacterial byproducts (35, 36). Therefore, clarifying the role of CPB2 toxin in IPEC-J2 cells can provide a scientific reference for preventing piglet diarrhea.

miRNAs play an important role in the resistance of piglets to infectious diarrhea caused by pathogenic bacteria. Bao et al. (37) found that miR-331-3p regulated Salmonella infection in pigs through the expression of the target protein VAV2. miR-221 targeted *FOS*, miR-125b targeted *MAPK14* and miR-27b targeted *IFNG* to regulate Salmonella infection in pigs (38). But there is little information on miR-132 in this area. Studies have found that miR-132 is closely related to inflammation. Alzahrani et al. (39) indicated that enhancing miR-132 expression by aryl hydrocarbon receptor attenuated tumorigenesis associated with chronic colitis. Yang et al. (40) demonstrated that miR-132 was down-regulated in LPS-induced inflammation, and targeting *TRAF6* attenuated inflammatory damage. Zhang et al. (41) found that miR-132/212 targeted *SIRT1* to activate NLRP3 inflammasome to promote LPS-induced apoptosis.But its function in piglet diarrhea is rarely studied. In our study, ssc-miR-132 was significantly down-regulated in IPEC-J2 cells treated with rCPB2 toxin. This suggested that it plays a role in proliferation, apoptosis and inflammation caused by rCPB2 toxin

The LDH is used as a sign of cell membrane damage and toxicity. Under normal circumstances, the release of LDH is very small. When the cell membrane permeability increases, the release of LDH increases(42, 43).In this study, we found that after IPEC-J2 cells were exposed to rCPB2 toxin, the LDH content increased. Overexpression of ssc-miR-132 reversed this result and decreased release of LDH. This result indicated that overexpression of ssc-miR-132 can alleviate the toxic damage of rCPB2 to IPEC-J2 cell membranes. Inflammatory cytokines are indicators to measure inflammation. *TNF-α* is a pro-inflammatory cytokine. It can activate NF-κB and induce the release of *IL-6*, *IL-8* and other pro-inflammatory factors, thereby increasing the impact on the intestinal mucosal barrier damage (44). *IL-6* is a multifunctional inflammatory factor, which mainly plays a role in the immune response of cells and body fluids, and has the functions of regulating immunity and inducing inflammation (45). *IL-8* is increased in inflammation and can cause intestinal mucosal damage (46). Current experiments showed that the rCPB2 toxin increased pro-inflammatory cytokines (*TNF-α*, *IL-6* and *IL-8*), while ssc-miR-132 mimics weakened their expression. This indicated that ssc-miR-132 can reduce the inflammatory damage caused by rCPB2. Further experiments indicated that overexpression of ssc-miR-132 overturned the inhibition of IPEC-J2 cell viability and promoted the expression of anti-apoptotic protein Bcl-2 and inhibited the expression of pro-apoptotic proteins Caspase-3 and Bax, thereby inhibiting the apoptosis induced by rCPB2 toxin.

Dachshund family transcription factor 1 (*DACH1*) is a key factor in determining cell fate (47). Studies have indicated that *DACH1* is necessary in the occurrence of certain inflammations (48, 49). *DACH1* is closely related to the cytokine *IL-8* and inhibits transcription of AP-1 and NF-κB by DS domain, and reduces *IL-8* expression and promoter activity (50). We found that knockdown of *DACH1* reduced the release of *TNF-α, IL-6* and *IL-8*, and attenuated the inflammatory damage caused by rCPB2. Previous studies observed that *DACH1* inhibited cell proliferation, clone formation and epithelial-mesenchymal transition by regulating the expression of *cyclin D1*, *IL-8* and other signal molecules, and prevents cell cycle progression (51–53). The results of this study are similar. Our datas showed that down-regulation of *DACH1* promoted cell proliferation and attenuated apoptosis. Through dual luciferase experiments, we proved that ssc-miR-132 and *DACH1* do have a targeting relationship, and the they have opposite effects. Finally, rescue experiments showed that *DACH1* attenuated the role of ssc-miR-132 in IPEC-J2 cells induced by rCPB2 toxin.

In summary, we demonstrated that ssc-miR-132 reduces the inflammatory injury and apoptosis induced by rCPB2 toxin in IPEC-J2 cells through negative regulation of *DACH1*. This is helpful for the treatment of enteritis and provides a scientific basis for the prevention and control of piglet diarrhea.

## Experimental procedures

### Sample collection

The seven-day-old healthy Landrace ×Yorkshire piglets were fed with *C. perfringens* type C bacteria liquid to establish an animal model (54). Normal and infected piglet duodenum, ileum, jejunum and other tissues were collected and immediately were placed in liquid nitrogen, and storage at −80 °C. All animal experiments are conducted under the guidance of the Ethical Committee of Experimental Animal Center of Gansu Agricultural University (approval number 2006-398).

### Cell culture

Human embryonic kidney-293T (HEK-293T) cells and IPEC-J2 cell line were purchased from BeNa Culture Collection (BNCC, Beijing, China). IPEC-J2 cells were cultured in DMEM/F12 (HyClone, Logan, UT, USA). The HEK-293T cells were cultured in DMEM (HyClone, Logan, UT, USA). DMEM/F12 and DMEM were supplemented with 10% FBS (Evergreen, Hangzhou, Zhejiang, China) and 1% penicillin(100 U/ml)/streptomycin (100 μg/ml) (Hyclone, Logan, UT, USA) at 37 °C in a humidified and 5% CO_2_ atmosphere.

### Cell transfection

miR-132 mimics, inhibitor and their respective negative controls (mimics NC and inhibitor NC) were purchased from RiboBio (Guangzhou, Guangdong, China). si-DACH1 (5’-GGCAGCUUCAACAGAUAGUTT-3’) and its negative control (si-NC) were synthesized by GenePharma (Shanghai, China). DACH1 3’UTR mutation vector and pcDNA3.1-DACH1 were synthesized by GENEWIZ (Suzhou, Jiangsu, China). When cells reached 50%-60% confluence, Lipofectamine™ 2000 reagent (Invitrogen, Carlsbad, CA, USA) was used to import the plasmids above into HEK-293T and IPEC-J2 cells as demanded.

### Lactate dehydrogenase (LDH) cytotoxicity assay

The cytotoxicity of rCPB2-treated cells was evaluated by LDH cytotoxicity assay kit (Jiancheng, Nanjing, Jiangsu, China). IPEC-J2 cells were incubated in 24 well plates (2 × 10^4^ cells/well) and transfected, then treated with rCPB2 (20 μg/mL) for 24 h. The toxin recombination method and concentration selection have been described in the previous paper (55). After 24 h, cells were collected for cytotoxicity detection according to the instructions.

### CCK8 assay

IPEC-J2 cells were seeded in 96‐well plates (3 × 10^3^ cells/well) and cultured 24 h, and they were transfected and treated with rCPB2 for 24 h. Add 10 μl CCK8 (Beyotime, Shanghai, China) solution into each well. After 2 h, absorbance was measured at 450 nm using a multimode microplate reader (Thermo Fisher Scientific, Waltham, MA, USA).

### 5-Ethynyl-2-deoxyuridine (EdU) staining

After IPEC-J2 cells were cultured in 24-well dishes (2 × 10^4^ cells/well) for 24 h, miR-132 mimics (100 nM), inhibitor (300 nM), pcDNA3.1-DACH1 (1 μg), si-DACH1 (300 nM) and their respective negative controls were transfected into cells. And rCPB2 toxin was used to treat cells after 24 h. Then, cells were stained using EdU cell proliferation kit (Beyotime, Shanghai, China) according to the instructions.

### TUNEL staining

The TUNEL kit (Beyotime, Shanghai, China) was used to detect cell apoptosis. The transfected cells were treated with 20 μg/mL rCPB2 for 24 h. Then cells were dyed according to the instructions, and finally observed under the microscope.

### qRT-PCR

Total RNA was extracted from IPEC-J2 cells using Trizol reagent (TransGen Biotech, Beijing, China). Its concentration and purity were measured at 260 nm and 280 nm. *Evo M-MLV* RT Kit with gDNA Clean for qPCR (Accurate Biotech, Changsha, Hunan, China) and Mir-X™ miRNA First-Strand Synthesis Kit (Takara, Dalian, Liaoning, China) were applied for the transcription of RNA into complementary DNA (cDNA). Quantitative reverse transcription-polymerase chain reaction (qRT-PCR) analysis was conducted with the application of SYBR Green Premix Pro Taq HS qPCR Kit (Accurate Biotech, Hunan, China). *GAPDH* and *U6* were used as normalized control for mRNAs and miRNAs. The gene expression value was calculated using the 2 ^−(ΔΔCt)^ method (56). The primers are shown in table 1.

**Table 1.**
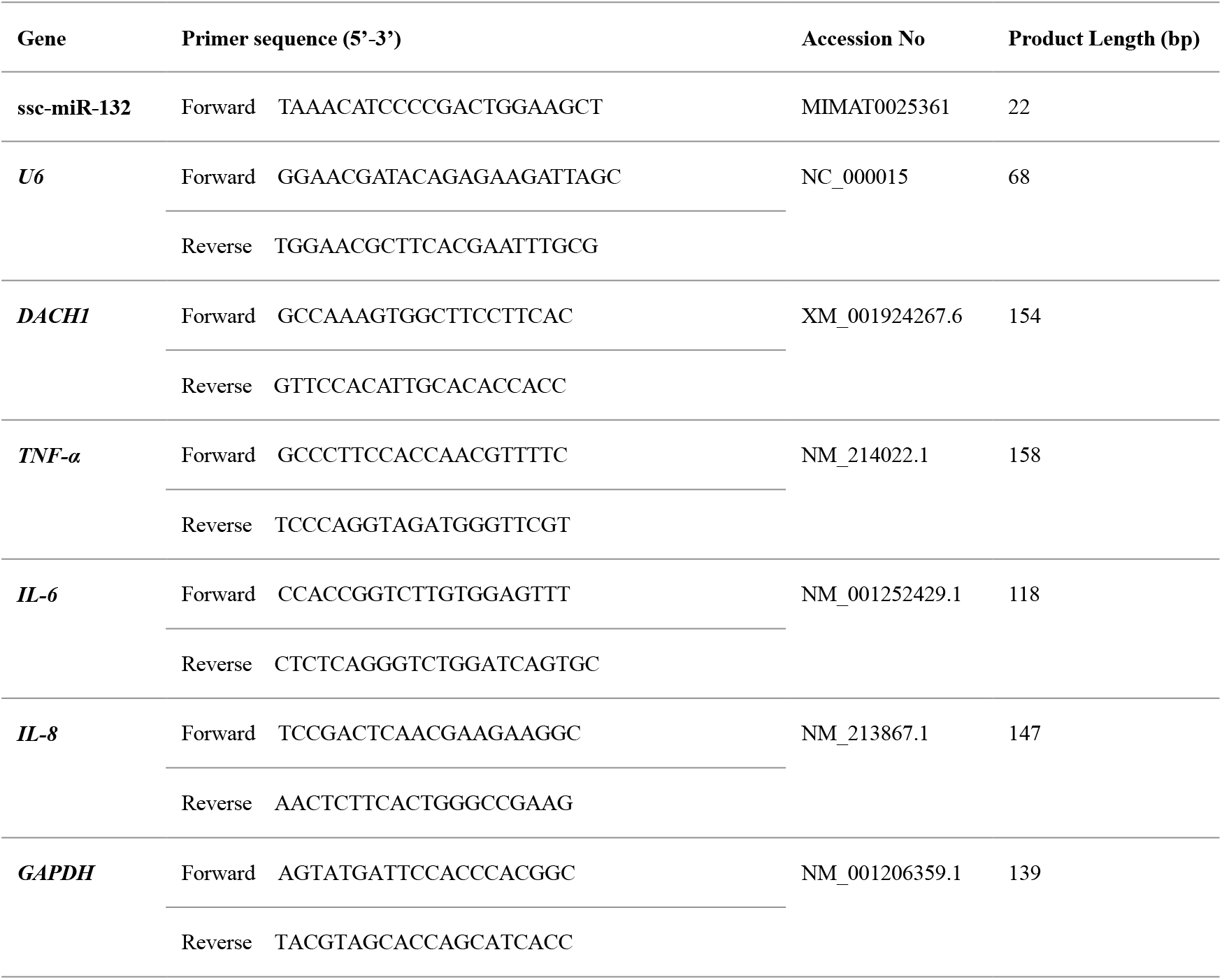
Primers used for qRT-PCR analysis

### Protein extraction and Western blot analysis

After cells were placed in a 6-well plate (3 × 10^5^ cells per well), the cells were transfected and treated with rCPB2 for 24 h. Cells were lysed with RIPA Lysis Buffer (Beyotime, Shanghai, China) to extract protein. Protein content was detected with BCA Protein Assay Kit (Beyotime, Shanghai, China). Then, the protein was electrophoresed by 12% SDS-PAGE (Solarbio, Beijing, China) for 90 min and was transferred to PVDF membrane (Beyotime, Shanghai, China) for 60 min. Afetr blocking with 5% skimmed milk (Beyotime, Shanghai, China) for 1 h, membranes were incubated at 4 °C overnight with primary antibody (bs-4222R, Bioss, 1:1000). Then they were washed three times in Tris-buffered saline with Tween (TBST) at room temperature in a decolorizing shaker after incubation for 5min each time. In addition, horseradish peroxidase (HRP)-labeled secondary antibody (bs-0295G-HRP, Bioss, 1:3000) incubated at room temperature for 30 min, and washed with TBST on a decolorizing shaker three times at room temperature for 5 min each time. Finally, an enhanced chemiluminescence (ECL) kit (NCM Biotech, Suzhou, China) were used to detect protein bands.

### Bioinformatics analysis

PicTar (https://pictar.mdc-berlin.de/), miRDB (http://mirdb.org/) and TargetScan (http://www.targetscan.org/vert_72/) were used to predict the target genes of ssc-miR-132. The miRNA mature sequences were downloaded from miRBase (http://www.mirbase.org/). The 3’-UTR sequence of the target gene was downloaded from the Ensembl genome browser (http://asia.ensembl.org/index.html).

### Dual luciferase reporter assay

DACH1 3’-UTR fragments containing the putative wild-type (WT) and mutant (MUT) were inserted into a dual-luciferase expression vector pmirGLO (Thermo Fisher Scientific, Waltham, MA, USA) with the *Xho*I and *Sal*I restriction endonucleases (Takara, Dalian, Liaoning, China) to produce the plasmid pmirGLO-DACH1 3’UTR-WT and pmirGLO-DACH1 3’UTR-Mut, respectively. Subsequently, each recombinant plasmid was cotransfected with ssc-miR‐132 mimic or mimics NC into 293T cells in 24-well plates using Lipofectamine™ 2000 reagent according to the manufacturer’s instruction. After 48 h, the Dual Luciferase Reporter Assay System (Promega, Madison, WI, USA) was employed to measure the activities of renilla and firefly luciferases in each group. Data was evaluated by normalizing the firefly luciferase activity to the renilla luciferase activity.

### Statistical analysis

All data were presented as mean ± SD and was performed using GraphPad Prism 6 software, and the two groups were compared using unpaired two-tailed Student’s t-tests. *P*-value < 0.05 was regarded as statistically significant.

## Acknowledgements

This research was funded by the Discipline Construction Fund Project of Gansu Agricultural University (GSAU-XKJS-2018-042).

## Conflict of interest

The authors declare that they have no conflicts of interest with the contents of this article.

## Abbreviations

*C. perfringens* type C: *Clostridium perfringens* type C
IPEC-J2: intestinal porcine epithelial
CPB2: *Clostridium perfringens* beta2
rCPB2: recombinant *Clostridium perfringens* beta2
LDH: lactate dehydrogenase
*TNF-α*: tumor necrosis factor
*IL-6*: interleukin-6
*IL-8*: interleukin-8
*DACH1*: dachshund family transcription factor 1
miRNAs: MicroRNAs
HEK-293T: Human embryonic kidney-293T
EdU: 5-Ethynyl-2-deoxyuridine
SDS-PAGE: odium dodecyl sulfate polyacrylamide gel
PVDF: polyvinylidene fluoride
IS: susceptible group

